# Genetic interactions drive heterogeneity in causal variant effect sizes for gene expression and complex traits

**DOI:** 10.1101/2021.12.06.471235

**Authors:** Roshni A. Patel, Shaila A. Musharoff, Jeffrey P. Spence, Harold Pimentel, Catherine Tcheandjieu, Hakhamanesh Mostafavi, Nasa Sinnott-Armstrong, Shoa L. Clarke, Courtney J. Smith, VA Million Veteran Program, Peter P. Durda, Kent D. Taylor, Russell Tracy, Yongmei Liu, Craig W. Johnson, Francois Aguet, Kristin G. Ardlie, Stacey Gabriel, Josh Smith, Deborah A. Nickerson, Stephen S. Rich, Jerome I. Rotter, Philip S. Tsao, Themistocles L. Assimes, Jonathan K. Pritchard

## Abstract

Despite the growing number of genome-wide association studies (GWAS), it remains unclear to what extent gene-by-gene and gene-by-environment interactions influence complex traits in humans. The magnitude of genetic interactions in complex traits has been difficult to quantify because GWAS are generally underpowered to detect individual interactions of small effect. Here, we develop a method to test for genetic interactions that aggregates information across all trait-associated loci. Specifically, we test whether SNPs in regions of European ancestry shared between European American and admixed African American individuals have the same causal effect sizes. We hypothesize that in African Americans, the presence of genetic interactions will drive the causal effect sizes of SNPs in regions of European ancestry to be more similar to those of SNPs in regions of African ancestry. We apply our method to two traits: gene expression in 296 African Americans and 482 European Americans in the Multi-Ethnic Study of Atherosclerosis (MESA) and low-density lipoprotein cholesterol (LDL-C) in 74K African Americans and 296K European Americans in the Million Veteran Program (MVP). We find significant evidence for genetic interactions in our analysis of gene expression; for LDL-C, we observe a similar point estimate although this is not significant, likely due to lower statistical power. These results suggest that gene-by-gene or gene-by-environment interactions modify the effect sizes of causal variants in human complex traits.

## Introduction

Over the last two decades, genome-wide association studies (GWAS) have demonstrated that human complex traits are influenced by many thousands of causal variants, each with small additive effects. What remains unclear is the extent to which traits are influenced by interactions between these variants, or between variants and the environment. Despite the dramatic increases in study size, GWAS are underpowered to detect individual gene-by-gene interactions of small effect. Testing for gene-by-environment interactions is similarly difficult, but with the added complication that the “environment” is notoriously hard to quantify. Thus, even though a handful of large-effect interactions have been identified^1–6^, the overall role of genetic interactions in complex trait architecture is yet to be determined.

Here, we test for genetic interactions by assessing whether causal variant effect sizes differ between populations. We use population differences in causal effect sizes as a proxy for genetic interactions because self-reported descriptors of population identity often loosely correlate with both genetic variation and environmental factors^7^. For example, in the United States, self-reported race often correlates with environmental exposures such as access to healthcare, due to a historical legacy of structural racism that extends into the present day^8^. This drives substantial environmental differences between populations, and if two populations have sufficiently different environmental backgrounds, then the existence of gene-by-environment interactions can produce modest differences in causal variant effect sizes. The existence of differential gene-by-gene interactions between populations would likewise produce differences in causal variant effect sizes.

However, comparing causal variant effect sizes between populations is rife with challenges. The causal variants underlying human complex traits are generally unknown and instead, GWAS typically identify single nucleotide polymorphisms (SNPs) that are statistically associated with the trait due to strong linkage disequilibrium (LD) with the causal variant(s). Due to differences in LD structure, these trait-associated SNPs may not be equally correlated with the same causal variant in two different populations, resulting in different marginal effect sizes. This is especially true if the causal variant is private, or only present in a single population. Thus, although several studies have observed differences between populations in the marginal effect sizes of trait-associated SNPs^9–12^, this could correspond both to differences in the effect sizes of causal variants themselves and to differences in LD structure.

These questions have been addressed further with statistical methods that leverage LD reference panels to account for differences in LD structure between populations^13,14^. These studies have found modest differences in causal variant effect sizes for both gene expression and complex traits. However, these existing methods are limited by their reliance on accurate LD reference panels and their difficulty in accounting for rare or population-specific causal variants. Furthermore, these methods are not suitable for application to recently admixed populations such as African Americans and Latin Americans due to the complexities of long-range admixture LD.

In this paper, we compare the genetic architecture of gene expression and low-density lipoprotein cholesterol (LDL-C) between African Americans and European Americans. Using data from the Multi-Ethnic Study of Atherosclerosis (MESA) and the Million Veteran Program (MVP), we first compare the marginal effect sizes of trait-associated SNPs when estimated from European Americans and from African Americans. We next quantify the contribution of local and global ancestry to phenotypic variance. Lastly, we leverage the multiple ancestries in the genomes of admixed populations to test for the existence of genetic interactions. Admixed African American genomes contain regions of European ancestry that share the same local LD structure as the genomes of European Americans. Within these regions of shared ancestry, we can compare variant effect sizes between populations without bias from differences in LD structure. Specifically, we hypothesize that in the absence of gene-by-gene or gene-by-environment interactions, SNPs will have the same effect sizes in European Americans and regions of European ancestry in African Americans. Conversely, we hypothesize that the presence of genetic interactions will drive the causal effect sizes of SNPs in regions of European ancestry in African Americans to be more similar to those of SNPs in regions of African ancestry.

## Material and Methods

### Genotype and phenotype datasets

#### Multi-Ethnic Study of Atherosclerosis (MESA)

For MESA, we obtained phased whole genome sequencing data and gene expression data in peripheral blood mononuclear cells (PBMCs) from TOPMed Freeze 8. After filtering individuals based on ancestry, as we describe below, the MESA dataset comprised 296 individuals who self-reported race as Black or African American and 482 individuals who self-reported race as White. We henceforth use the term “African American” to refer to all individuals who self-report race as Black or African American. Analogously, we use the term “European American” to refer to all individuals who self-report race as White and cluster with individuals of European ancestry in principal components analysis of genotypes.

380 of these individuals had gene expression data available at two exams, spaced five years apart. For these individuals, we selected the time of exam to use such that the proportions of certain covariates (sex, time of exam, sequencing center) were approximately balanced between European Americans and African Americans. Briefly, this was done by iterating through this set of individuals ten times and changing the time of exam used for that individual if doing so would increase the similarity of covariate proportions between the two populations.

As done previously by^15^, gene-level expression quantification was based on the GENCODE 26 annotation, collapsed to a single transcript model for each gene using a custom isoform collapsing procedure. Gene-level read counts were obtained with RNA-SeQC v1.1.9^16^. We selected genes with expression thresholds of >0.1 TPM in at least 20% of samples and ≥6 reads in at least 20% of samples, thresholding separately for European Americans and African Americans in both cases. A total of 10,870 genes passed this filtering step. We log-transformed gene expression measurements and used these transformed phenotypes in all downstream analyses. We selected biallelic SNPs with a MAF > 0.05 and minor allele sample count > 5 in both European Americans and African Americans.

#### Million Veteran Program (MVP)

For MVP, we used GRCh37 genotype calls processed and subject to quality control as described in^17^. Data were imputed with IMPUTE using the 1000 Genomes Phase 3 reference panel^18,19^. As previously done for MVP^17^, the population of each individual (i.e. African American or European American) was determined by HARE^20^. Using KING coefficients^21^, we removed relatives who were closer than 3rd degree cousins, which left 73,788 African American and 296,124 European American individuals. For all analyses, we used the maximum LDL-C measurement for each individual across all time points. In addition, we numerically adjusted LDL-C measurements for statin usage by multiplying measurements by 0.7 if an individual was inferred to be on statin medication. We inferred that individuals were on statin medication if a statin prescription was filled within the length of the prescription plus a buffer of 15 days within the LDL-C measurement date.

### Inferring global and local ancestry

We inferred global ancestry for admixed African American individuals with supervised ADMIXTURE using default program parameters^22^. We used 99 CEU individuals and 108 YRI individuals from 1000 Genomes Phase 3 as our reference populations. We filtered for biallelic SNPs with MAF > 0.05 in both the admixed population and the reference populations, and again filtered for MAF > 0.1 after merging the admixed and reference datasets. We pruned SNPs with an *r*^2^ value > 0.1.

We inferred local ancestry with RFMix v1.5.4, using no EM iterations and default program parameters^23^. We assumed 8 generations since the time of admixture between an African population and a European population^24^. We again used 99 CEU individuals and 108 YRI individuals from 1000 Genomes Phase 3 as our reference populations. We used biallelic SNPs with MAF > 0.05 in both the admixed population and the reference populations, and removed SNPs with an *r*^2^ value > 0.5.

In both datasets, we excluded African Americans with < 0.5 global African ancestry from downstream analyses. We also excluded one European American individual from MESA who did not cluster with individuals of European ancestry in principal components analysis of genotypes.

### Comparing marginal SNP effect sizes between populations

#### Gene expression (MESA)

To identify SNPs affecting expression in *cis*, we filtered for SNPs within 100 kb of the TSS for each gene. We ascertained trait-associated SNPs in a randomly sampled subset of 232 European Americans using ordinary least squares. This regression included ten covariates that were significantly correlated with expression phenotypes: sequencing center; time of exam; sex; genotype PC 2, which captures structure within European Americans; and six covariates corresponding to a one-hot encoding of recruitment site (Figure S2, Figure S3A).

For each gene, we focused on the most significant SNP and ascertained significant SNP-gene associations by applying a false discovery rate of 0.01 to correct for multiple testing, as done by^15^. All downstream analyses were performed on these significant associations. Furthermore, all downstream analyses excluded the individuals who were used to ascertain trait-associated SNPs.

For each significant SNP-gene association, we performed two separate regressions to estimate *β*_*AA*_, the effect size in African Americans, and *β*_*EA*_, the effect size in European Americans, respectively. For each regression, we again included covariates significantly correlated with expression phenotypes. To estimate *β*_*EA*_, we used sequencing center, time of exam, sex, genotype PC 2, and recruitment site as above. To estimate *β*_*AA*_, we used sequencing center, time of exam, sex, recruitment site, and global African ancestry fraction. (We did not include genotype PC 1 as a covariate despite its significant association with expression because this is highly correlated with global African ancestry fraction (Figure S3B).) We estimated *β*_*EA*_ in 250 European Americans and randomly sampled an equal number of African Americans to estimate *β*_*AA*_.

#### LDL-C (MVP)

We ascertained genome-wide significant SNPs in 318,953 UK Biobank White British individuals. After applying genomic filters (MAF ≥ 0.01, missing genotype rate *≤* 0.05, Hardy-Weinberg equilibrium with a cutoff of *p* < 1 × 10^−50^), we tested for association with inverse-variance quantile normalized phenotypes using a linear model (–glm) in plink with the covariates age, sex, assessment center, and statin usage. Significant variants (*p* < 5 × 10^−8^) were clumped and thinned to leave at most one independent SNP per 0.1 cM^25^.

To estimate effect sizes of these variants in MVP, we extracted variants from the imputed genotype set using 1000 Genomes Phase 3 as our reference panel. We filtered for MAF ≥ 0.003 in European Americans and African Americans, leaving 122 independent SNPs. Our covariates included age, sex, global ancestry, and genotype PC 1, which stratifies European Americans and is the only principal component associated with LDL-C after residualizing on the other covariates. Principal components were calculated on all individuals in the MVP dataset with HARE^17^.

To estimate effect sizes from the 74K African Americans (*β*_*AA*_), we used linear regression in plink (–glm) and included the covariates above. We likewise randomly sampled an equal number of European Americans and estimated effect sizes (*β*_*EA*_).

#### Comparison of effect sizes

We used total least squares (TLS) regression to assess the slope of the relationship between 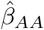 and 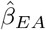 Estimates of SNP effect sizes are statistically noisy, and unlike ordinary least squares, total least squares is robust to uncertainty in the x-axis variable. Because we used the same number of samples to estimate *β*_*AA*_ and *β*_*EA*_, their standard errors will be comparable, as is necessary for TLS regression. We created 1000 bootstrap replicates for each trait by sampling with replacement over SNPs and report the 95% confidence interval (CI) of the slope as defined by the 0.025 and 0.975 quantiles.

### Quantifying role of ancestry in phenotypic variance

We constructed a series of phenotypic models and compared the proportion of phenotypic variance explained by each model. We fit each model in a training set comprising 80% of the data (for gene expression, 237 African Americans and 200 European Americans; for LDL-C, 52K African Americans and 52K European Americans). We computed the proportion of variance explained as 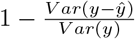 in a test set comprising the remaining 20% of the data (for gene expression, 59 African Americans and 50 European Americans; for LDL-C, 22K African Americans and 22K European Americans). This quantity can be interpreted as measuring the decrease in residual variance relative to phenotypic variance. For gene expression, we report the average variance explained across all significant genes.

We note that this procedure differs from our previous analysis in two ways. First, we fit the models below by performing a regression on the joint sample of African Americans and European Americans, while previously, we performed a regression in each population separately. Second, though we previously downsampled the number of African Americans in MESA, here we included all 296 African Americans to maximize our power to estimate local ancestry-specific effect sizes. (In addition, because African American genomes contain both African and European ancestry, it is not as useful to downsample the number of African Americans for these analyses.)

We first modeled the phenotype *y* in an individual *i* with only technical covariates (*c*). For gene expression, this consisted of sex and batch (sequencing center, time of exam, and recruitment site); for LDL-C, this consisted of age and sex.

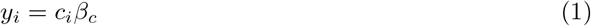

Consecutive models added an indicator variable for race (*r*), followed by genome-wide descriptors of ancestry (*θ*). Specifically, *θ* includes global African ancestry fraction and genotype principal components that stratify European Americans (PC 2 for gene expression, see Figure S3; and PC 1 for LDL-C^17^).

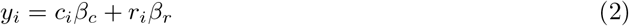

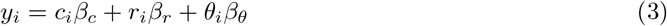

We next included a local ancestry covariate (*γ*) that measures the number of haplotypes with African ancestry at the trait-associated SNP. For gene expression, we averaged across all SNP-gene associations to report the variance explained by local ancestry. On the other hand, for LDL-C, we summed across all trait-associated SNPs to report the variance explained.

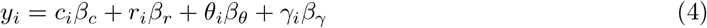

Lastly, we included the genotype at trait-associated SNPs. We modeled the genotype with ancestry-specific effect sizes, given that differences in LD structure produce differences in the marginal effect sizes of trait-associated SNPs. Rather than adding a single term for trait-associated SNPs (e.g. *g*_*i*_*β*_*g*_), we added two terms, *g*_*i,A*_*β*_*A*_ and *g*_*i,E*_*β*_*E*_. We define *g*_*i,A*_ as the number of alternate alleles with African local ancestry and *g*_*i,E*_ as the number of alternate alleles with European local ancestry. *g*_*i,A*_ and *g*_*i,E*_ therefore sum to *g*_*i*_, the total genotype, and *β*_*A*_ is the effect size in African local ancestry while *β*_*E*_ is the effect size in European local ancestry. Once again, to report the variance explained, we averaged across all SNP-gene associations for gene expression and summed across all trait-associated SNPs for LDL-C.

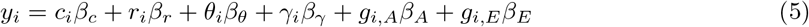

### Testing for genetic interactions

#### Overview of model

We constructed a phenotypic model in which we introduce the parameter *δ* to measure differences in the marginal effect size of trait-associated SNPs in regions of European ancestry in African Americans compared to European Americans.

We extend Equation 5, modeling the phenotype *y* for a single individual *i* as follows:

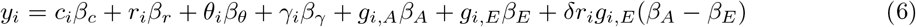

As described above, the first four terms (*c*_*i*_, *r*_*i*_, *θ*_*i*_, *γ*_*i*_) are technical covariates; race; global ancestry and principal components; and local ancestry, respectively. The next two terms (*g*_*i,A*_*β*_*A*_, *g*_*i,E*_*β*_*E*_), model ancestry-specific effect sizes of trait-associated SNPs.

In the final term, we introduce the parameter *δ*, which measures the extent to which marginal effect sizes of SNPs in regions of European ancestry in African Americans differ from those in European Americans. Using the parameter *δ*, we can indirectly test whether causal variant effect sizes differ between African Americans and European Americans. When *δ* equals 0, the marginal effect size of a SNP in a region of European ancestry in an African American is equal to *β*_*E*_; as *δ* approaches 1, the marginal effect size approaches *β*_*A*_. Thus, under the null hypothesis that causal variant effect sizes are identical between populations, *δ* will be equal to 0. However, if causal variant effect sizes differ between populations because they are modified by the genome and/or environment, *δ* will be greater than 0. (We note that a value of *δ* equal to 0 is not evidence for the absence of any genetic interactions; rather, it indicates that genetic interactions do not differ enough between populations to produce differences in causal variant effect sizes.)

#### Fitting the model

To fit this model, we began by initializing 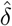 to a random value on the interval [0, 1], which is the most biologically intuitive range of values for *δ* (see Figure 3A). We next optimized 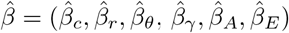 conditional on this value of 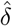, and we then optimized 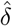 conditional on 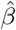. For both gene expression and LDL-C, we performed this regression marginally on each SNP. In other words, conditional on 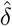, we estimated 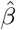 for each SNP independently of the rest. We continued this iterative optimizing with ordinary least squares regression until 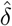 converged (i.e. did not change by >.0001). Though 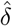 was initialized on the interval [0, 1], the optimization procedure itself was unconstrained. Additionally, we found that regardless of the initial value of 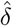, our optimization procedure converged to the same value. The optimization method converged quickly for both datasets (22 iterations for gene expression, 18 for LDL-C). For the gene expression data, we estimated one value of *δ* from all SNP-gene associations to avoid overparameterization. For the LDL-C data, we estimated one value of *δ* across all trait-associated SNPs. To construct 95% confidence intervals for 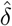, we bootstrapped over SNPs and reported the 0.025 and 0.975 quantiles. (For gene expression, this procedure is equivalent to bootstrapping over genes because each gene is modeled by exactly one SNP.) We concluded that causal variant effect sizes are significantly different if the 95% CI does not include 0. To generate a likelihood surface for *δ*, we computed the log-likelihood of the data conditional on values of *δ* ranging from 0 to 1, with a step size of 0.01.

#### Assessing properties of the estimator 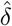

We first assessed the bias of our estimator 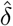 with simulations designed to emulate our analyses of gene expression in MESA. We simulated genotypes and phenotypes for 100 independent loci in 320 admixed African Americans and 500 Europeans. For each African American, we simulated global African ancestry fraction from a beta distribution (*α* = 7.9, *β* = 2.1) resembling the empirical distribution of global ancestry. For each locus, we simulated local ancestry conditional on global ancestry from a binomial distribution.

We then simulated the respective numbers of African and European genomes using the two-population out-of-Africa model as implemented in stdpopsim^26–29^. For each locus, we simulated 1% of chromosome 22 and filtered for SNPs that had MAF > 0.05 in both African and European genomes. To mimic ascertainment in a European population, we held out genomes for 250 European individuals; to mimic ascertainment in an African population, we held out genomes for two-thirds of all individuals who had two copies of African ancestry (i.e. *γ* = 2). We simulated causal and tag SNPs by jointly sampling at random from the set of all pairs of SNPs with *r*^2^ greater than a specified threshold in the ascertainment individuals. We conducted simulations with *r*^2^ thresholds of 0.6 and 0.8.

We simulated causal variant effect sizes from a bivariate normal distribution with a correlation of 0.85, which allowed causal variant effect sizes in African and European ancestries to differ (e.g. due to gene-by-gene or gene-by-environment interactions). We simulated phenotypes from causal SNP genotypes using the generative model we specified in Equation 6, ignoring the role of technical, race, and ancestry covariates. For simulations in which causal variant effect sizes differ between populations, we simulated five values of *δ* ranging between 0 and 0.8. We estimated *δ* by applying our iterative optimization procedure to the simulated phenotypes and tag SNP genotypes for all 100 loci. For each combination of hyperparameters (ascertainment population, *r*^2^ threshold, and simulated value of *δ*), we performed 10 simulations.

Lastly, we assessed the behavior of our estimator 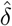 in the case where causal variant effect sizes are identical between populations. In principle, if the true marginal effect sizes *β*_*A*_ and *β*_*E*_ are identical, then the parameter *δ* is not identifiable. In practice, we do not expect the marginal effect sizes *β*_*A*_ and *β*_*E*_ to be identical due to differences in LD structure between African and European ancestries. Nevertheless, we investigated this further in both simulations and empirical data. In simulations, we used a similar framework to that described above, but we used a univariate normal distribution to simulate causal variant effect sizes that were identical between populations. In empirical data, we modified our model such that we could use *δ* to compare effect sizes between two randomly sampled, independent subsets of European Americans. On average, individuals in these two subsets have the same race, global ancestry, local ancestry, and environment. Thus, we expect that causal variant effect sizes are identical between subsets even in the presence of gene-by-gene or gene-by-environment interactions. To modify our model, we first excluded any African Americans with European ancestry at trait-associated SNPs. This ensured that *β*_*E*_ was estimated only from European Americans at trait-associated SNPs, and that *β*_*A*_ was estimated only from African Americans with African ancestry on both haplotypes at trait-associated SNPs. Next, we assigned a randomly sampled subset of the European Americans as a validation set. For gene expression, this was 100 individuals, and for LDL-C, this was 74K individuals. We then replaced the race indicator in the last term of the model with a validation set indicator. With this particular modification of the model, our estimator 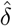 tests whether trait-associated SNPs have the same effect size in two randomly sampled, independent subsets of European Americans. If *δ* is estimated to be nonzero between these two subsets of European Americans, this would indicate that our estimator has pathological behavior in the case where causal effect sizes are identical between populations.

## Results

We performed analyses for gene expression and LDL-C, both of which are driven by a combination of genetic factors and environmental factors. We analyzed gene expression using MESA, a dataset with whole genome sequencing and bulk RNA-Seq in peripheral blood mononuclear cells for 296 African Americans and 482 European Americans. We analyzed LDL-C using MVP, a dataset with dense SNP genotyping and LDL-C measurements for 74K African Americans and 296K European Americans. Of existing human genetic datasets, MESA and MVP have some of the largest cohorts of admixed individuals for their respective phenotypes.

### Inferring global and local ancestry

We inferred global and local ancestry for the African American individuals in MESA and MVP. In both cases, we modeled African Americans as a two-way admixture between African and European populations that occurred 8 generations ago^24^. We estimated global ancestry using supervised ADMIXTURE with 1000 Genomes populations (CEU as European and YRI as African) as our reference populations^18,22^. The average global African ancestry of African American individuals is 0.80 in MESA and 0.82 in MVP, concordant with previous estimates from similar populations^30^ (Figure 1A). We performed local ancestry inference with RFMix using the same 1000 Genomes reference populations^23^. Global ancestry fractions from ADMIXTURE are highly correlated with those implied by RFMix (MESA *ρ* = 0.997, MVP *ρ* = 0.98) (Figure S1). As expected based on their admixture history, the local ancestry of African American individuals alternates between blocks of African and European ancestry along the genome and contains relatively large European blocks (mean length is 15 Mb in MESA, 14 Mb in MVP) (Figure 1B).

**Figure 1:**
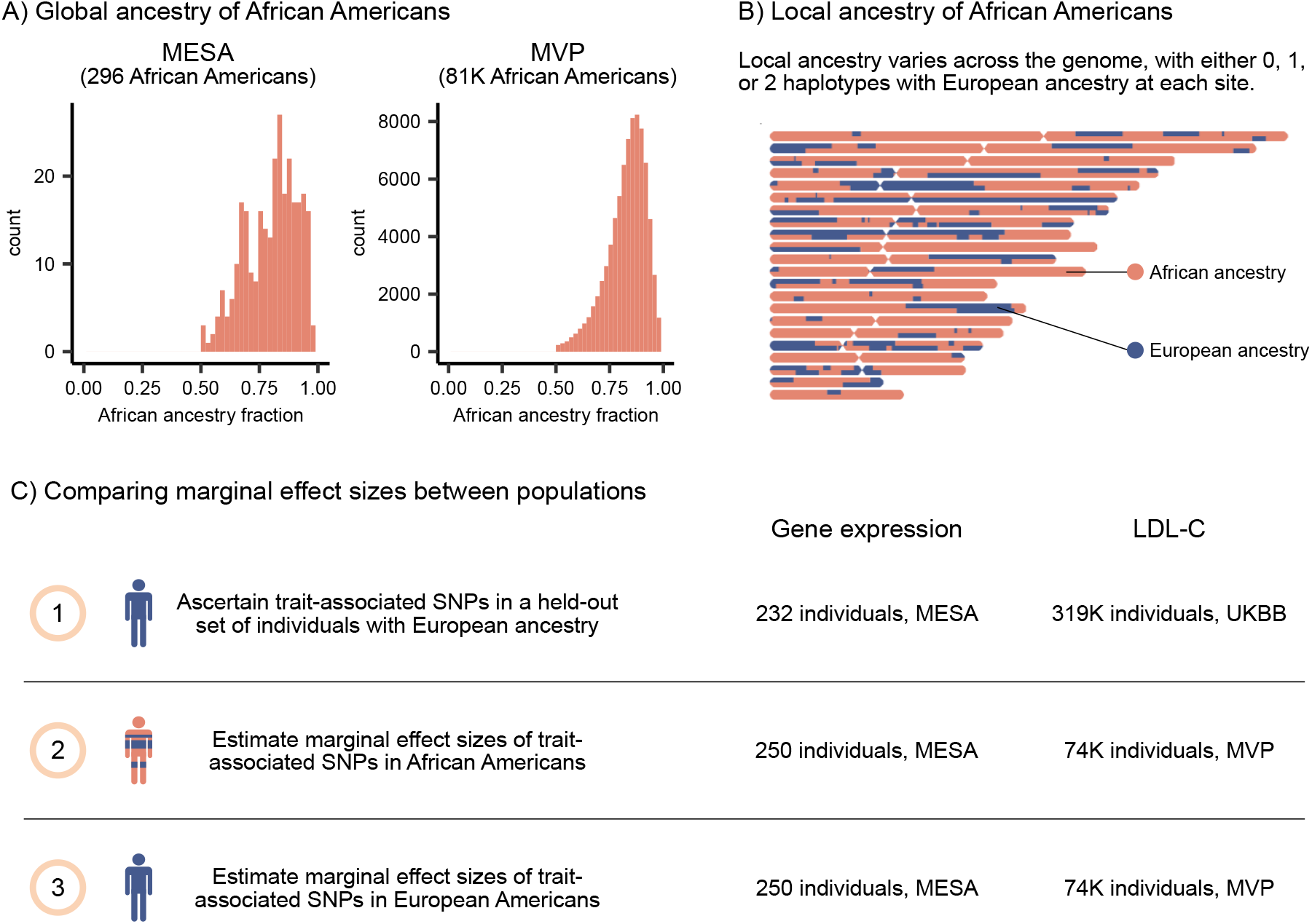
Schematic of the analysis pipeline. **A)** Global ancestry of African Americans is predominantly African, with an average global African ancestry fraction of 0.80 in MESA and 0.82 in MVP. **B)** Local ancestry for one sample individual in MESA. Individuals have either 0, 1, or 2 haplotypes with European ancestry at each position. **C)** We compare marginal effect sizes of SNPs between African Americans and European Americans.

### Comparing marginal SNP effect sizes between populations

We first sought to compare marginal effect sizes of trait-associated SNPs when estimated from European Americans and from African Americans (Figure 1C). We expect that marginal effect sizes of trait-associated SNPs will differ between the two populations due to known differences in LD structure between African and European ancestries, as well as potential differences in gene-by-gene or gene-by-environment interactions between the populations. However, the magnitude of this difference in marginal effect sizes is unclear. The observed magnitude of differences may be inflated by sampling error, particularly if one population has a small sample size. Additionally, effect sizes are usually largest in a discovery sample due to Winner’s Curse, which further exacerbates differences between discovery and replication datasets. To minimize these biases, we ascertained trait-associated SNPs in a held-out set of individuals and compared effect sizes in an equal number of African Americans and European Americans (Fig 1C).

We first log-transformed phenotype measurements for variance stabilization. We did not perform quantile normalization given that phenotypic variance might differ between populations^31^. We ascertained unlinked, trait-associated SNPs in individuals of European ancestry. For gene expression, we restricted our analyses to putative *cis*-acting variants (i.e. within 100 kb of TSS) because *cis*-acting variants have stronger effects than *trans*-acting variants and are more easily detected in modest sample sizes^15,32^. In the event that there were multiple SNPs associated with a gene, we chose the most significant SNP for downstream analyses^15^. We ascertained trait-associated SNPs (false discovery rate < 0.01) in a held-out subset of 232 European Americans in MESA, which resulted in 4,236 SNP-gene associations. For LDL-C, we ascertained trait-associated SNPs (*p* < 5 × 10^−8^) in 318,953 UK Biobank (UKBB) White British individuals, and clumped and thinned them, which resulted in 122 trait-associated SNPs. We performed all subsequent analyses on these trait-associated SNPs.

To compare marginal effect sizes between populations, we estimated the effect sizes of trait-associated SNPs separately in African Americans (*β*_*AA*_) and European Americans (*β*_*EA*_). For gene expression, *β*_*AA*_ and *β*_*EA*_ were each estimated from 250 individuals. For LDL-C, *β*_*AA*_ and *β*_*EA*_ were each estimated from 74K individuals. For each trait, we compared marginal effect sizes between the two populations by regressing effect sizes estimated from African Americans (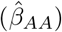) on effect sizes estimated from European Americans (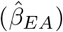) (Figure 2). We used total least squares (TLS) to perform the regression because it is robust to statistical noise in the independent variable (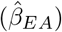), while ordinary least squares is not.

**Figure 2:**
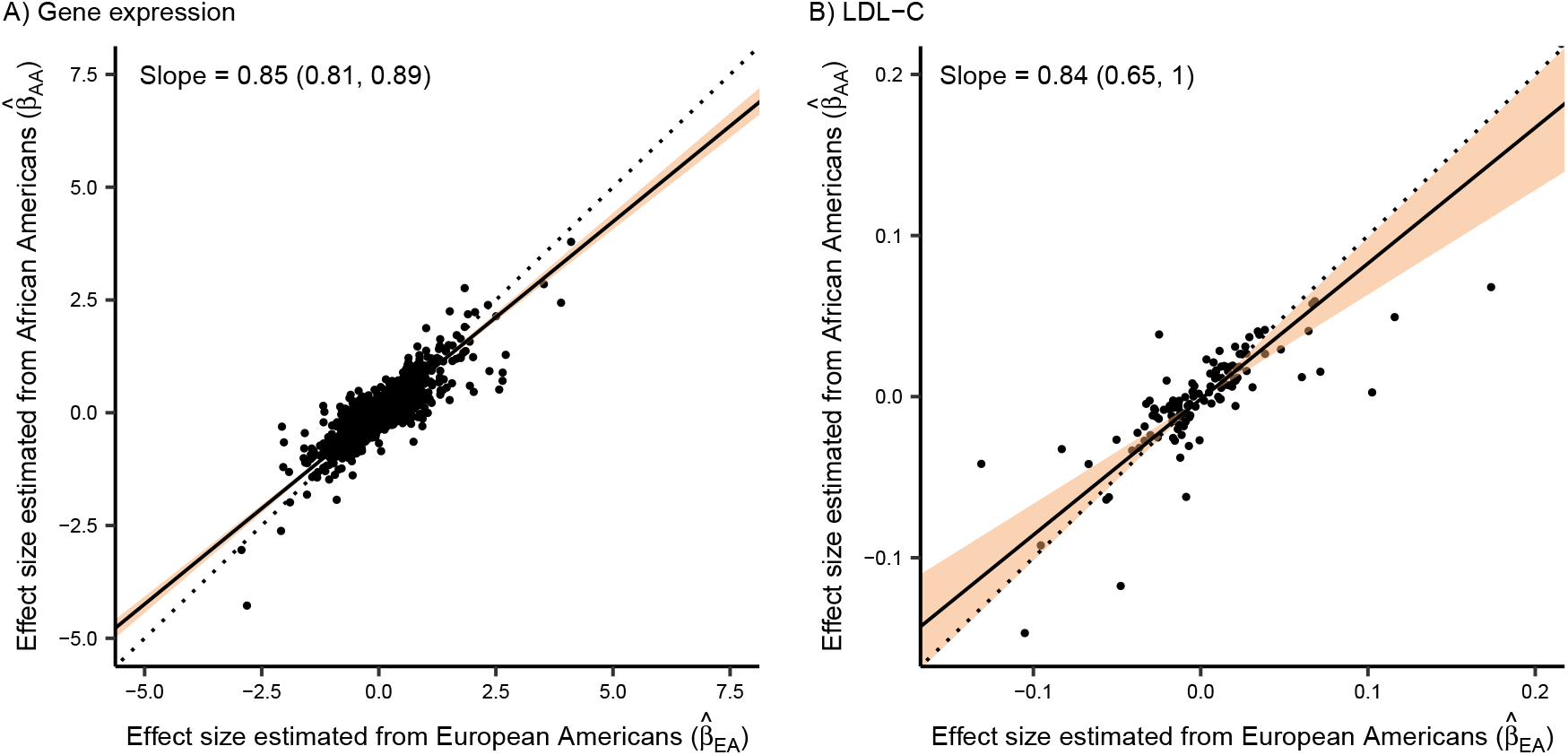
Comparing marginal SNP effect sizes between populations. We estimated effect sizes of trait-associated SNPs and regressed the effect size estimated from African Americans on the effect size estimated from European Americans. We represent the 95% bootstrap CI with the shaded region. Effect sizes estimated from African Americans are **A)** significantly smaller in magnitude than the corresponding effect sizes estimated from European Americans for gene expression and **B)** smaller but not significantly so for LDL-C.

For gene expression, effect sizes estimated from African Americans are significantly smaller in magnitude than the corresponding effect sizes estimated from European Americans, with a slope of 0.85 (95% CI of 0.81-0.89) (Figure 2). For LDL-C, we similarly observe a slope of 0.84, but this is not significantly different from 1 (95% CI of 0.65-1.01), likely due to the modest number of SNPs analyzed for this trait. Our observation that marginal effect sizes estimated from African Americans are smaller in magnitude can be at least partially explained by our ascertainment of trait-associated SNPs in individuals of European ancestry. Blocks of LD structure are smaller in populations of African ancestry than in populations of European ancestry, and the African Americans in MESA and MVP have a mean African global ancestry of approximately 80%. Thus, the correlation between causal variants and trait-associated SNPs ascertained in European populations will generally be weaker in African Americans than in European Americans, meaning that marginal effect sizes estimated from African Americans will have a smaller magnitude. Potential differences in gene-by-gene and gene-by-environment interactions between populations could also contribute to the observed differences in marginal effect sizes, but are unlikely to produce such a systematic shift in the magnitudes of effect sizes.

### Quantifying role of ancestry in phenotypic variance

Given that African Americans are admixed with both African and European ancestries, we next sought to assess the contribution of global and local ancestry to phenotypic variation. We quantified the contribution of both terms to phenotypic variation by constructing a series of phenotypic models and computing the amount of variance explained by each model. We fit each model to roughly 80% of our data allocated as a training set and computed the proportion of phenotypic variance explained by the model in a test set using the remaining 20% of our data. For gene expression, we report the average phenotypic variance explained across all genes.

We constructed five phenotypic models in total, where each model has an increasing number of terms relative to its predecessor. Our first phenotypic model (Table 1; Equation 1) included only technical covariates (sex and batch for gene expression; sex and age for LDL-C) and explains 18.41% of phenotypic variance for gene expression and 0.11% of phenotypic variance for LDL-C. Most of the variance explained by these covariates for gene expression is due to batch effects, as is common for RNA-Seq assays. We next added an indicator variable for race, which allows for race-specific phenotypic intercepts and can capture trait-relevant differences in environment between African American and European American populations^33^ (Equation 2). Compared to a model that only includes technical covariates, including race explains an additional 1.26% of variance in gene expression and 0.05% of variance in LDL-C.

**Table 1:** Quantifying role of ancestry in phenotypic variance. We constructed a series of linear models and computed the percentage of phenotypic variance explained. For both traits, we report the increase in the percentage of phenotypic variance explained by each model; for gene expression, we report the average increase across all genes. The variables in the models are defined as follows: *c*_*i*_ is a vector of technical covariates; *r*_*i*_ is a race indicator variable; *θ*_*i*_ is a vector of global African ancestry fraction and principal components; *γ*_*i*_ is a local ancestry covariate that measures the number of haplotypes with African ancestry at trait-associated SNPs; *g*_*i,A*_ is the number of alternate alleles with African local ancestry and *g*_*i,E*_ is the number of alternate alleles with European local ancestry

|  | Term added | Model | Additional variance explained (%) |  |
| --- | --- | --- | --- | --- |
|  |  |  | Gene expression | LDL-C |
| (1) | Technical covariates | $y_i = c_i\beta_c$ | 18.41 | 0.11 |
| (2) | Race | $y_i = c_i\beta_c + r_i\beta_r$ | 1.26 | 0.05 |
| (3) | Global ancestry & PCs | $y_i = c_i\beta_c + r_i\beta_r + \theta_i\beta_\theta$ | 0.00 | 0.01 |
| (4) | Local ancestry | $y_i = c_i\beta_c + r_i\beta_r + \theta_i\beta_\theta + \gamma_i\beta_\gamma$ | 0.03 | -0.04 |
| (5) | Genotype with ancestry-specific effect sizes | $y_i = c_i\beta_c + r_i\beta_r + \theta_i\beta_\theta + \gamma_i\beta_\gamma + g_{i,A}\beta_A + g_{i,E}\beta_E$ | 3.62 | 2.12 |

We next added global African ancestry fraction and genotype principal components to the model (Equation 3). These covariates can capture additional population structure: global African ancestry fraction stratifies African Americans, while the principal components we include stratify European Americans. In the context of gene expression, global ancestry and genotype principal components are known to be relevant for trait variation, potentially because they capture the effect of *trans* genetic variation on expression^2,1,33^. Surprisingly, we find that these terms have a small contribution to the overall phenotypic variance of both gene expression and LDL-C.

We next considered the importance of a local ancestry covariate that measures the number of haplotypes with African ancestry at each trait-associated SNP (Equation 4). Local ancestry could implicitly capture the effect of local genetic variation from SNPs that are not explicitly modeled; in the context of gene expression, these unmodeled, trait-associated SNPs are likely *cis*-acting variants. However, we find that including local ancestry does not explain much additional variance in either gene expression or LDL-C.

Lastly, we considered the role of trait-associated SNPs (Equation 5). Differences in LD structure between African and European ancestries result in different marginal effect sizes at trait-associated SNPs, as we see in Figure 2. Consequently, we modeled the genotype at trait-associated SNPs with ancestry-specific effect sizes. We find that trait-associated SNPs contribute considerably to trait variation, explaining an additional 3.62% of variance in gene expression and 2.12% of variance in LDL-C. Thus, we find that the genotype at trait-associated SNPs contributes substantially more to phenotypic variance than either local or global ancestry.

### Testing for genetic interactions

Finally, we looked for evidence of genetic interactions by testing whether causal variant effect sizes differ between populations. This is difficult to do with standard approaches due to the way in which LD structure can bias comparisons of marginal effect sizes. We therefore developed a model that leverages the multiple ancestries within admixed genomes to indirectly test whether causal variant effect sizes differ between populations. Specifically, we test whether a genetic variant in a region of European ancestry has the same marginal effect size in African Americans and European Americans. We assume that the regions of European ancestry in the African Americans and European Americans in our datasets are virtually identical with respect to LD structure, which means that differences in marginal effect sizes should reflect differences in causal effect sizes.

This assumption is based on the specific demographic history of African Americans and Europeans. Given the relatively short time since admixture in African Americans (approximately 8 generations), we expect that regions of European ancestry in modern-day African Americans feature the same LD structure as the European source population contributing to the admixture event^24^. Moreover, others have previously demonstrated that there is low Fst and high correlation of allele frequencies between various European populations^34–36^. Empirically, we also find that nearly all (95%) SNPs that are tightly linked (*r*^2^ > 0.8) in European Americans in MESA are also tightly linked in regions of European ancestry in African Americans in MESA. (In contrast, only 65% of SNPs that are tightly linked in European Americans are tightly linked in regions of African ancestry in African Americans.) Thus, we have extensive support for the assumption that the LD structure between trait-associated SNPs and causal variants is similar in European Americans and regions of European ancestry in African Americans.

Then, under the null hypothesis that genetic interactions do not impact causal variant effect sizes, causal variants will have an identical effect size in all populations, and trait-associated SNPs in regions of European ancestry will have the same marginal effect size in African Americans and European Americans. However, if genetic interactions drive differences in causal variant effect sizes between populations, trait-associated SNPs in regions of European ancestry will have different marginal effect sizes in African Americans and European Americans. Specifically, we hypothesize that in African Americans, the presence of genetic interactions will drive the marginal effect sizes of SNPs in regions of European ancestry to be more similar to those of SNPs in regions of African ancestry. As we noted previously, without accounting for LD structure, we would expect marginal effect sizes of trait-associated SNPs to differ between populations regardless of whether causal variant effect sizes do (i.e. regardless of whether genetic interactions exist). However, because we focus on regions of shared European ancestry in two different populations, our comparison of marginal effect sizes is not biased by differences in LD structure, nor by the possibility of private causal variants in European populations.

We test this hypothesis by developing a model that uses the parameter *δ* to measure the extent to which marginal effect sizes of SNPs in regions of European ancestry in African Americans deviate from those in European Americans (see Methods, Equation 6). Values of *δ* greater than 0 indicate that SNPs in regions of European ancestry in African Americans and European Americans have different marginal effect sizes. In addition, values of *δ* greater than 0 indicate that SNPs in regions of European ancestry in African Americans have effect sizes more similar to SNPs in regions of African ancestry in African Americans. Thus, values of *δ* greater than 0 provide evidence for a difference in causal variant effect sizes between populations.

For both traits, we fit this model to the trait-associated SNPs we previously ascertained. We expect that estimates of *δ* will be noisy at individual SNPs, so for each trait, we estimated a single shared value of *δ* across all SNPs. This results in one value of *δ* for gene expression, estimated from all SNP-gene associations, and one value for LDL-C, estimated from all LDL-associated SNPs. Because this model is non-linear, we iteratively optimized *δ* and all other coefficients, *β* = (*β*_*c*_, *β*_*r*_, *β*_*θ*_, *β*_*γ*_, *β*_*A*_, *β*_*E*_) with ordinary least squares until convergence. To construct a confidence interval for 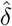, we bootstrapped over SNPs.

We first assessed the bias of our estimator 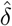. Using a standard demographic model, we simulated genotypes for admixed African Americans and Europeans. In order to simulate the LD structure present in our analyses of real data, we simulated phenotypes from causal SNP genotypes but estimated *δ* from tag SNP genotypes in simulations. We find that our estimates 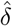 are well-correlated with the simulated values of *δ* regardless of ascertainment population (Figure S5). We next assessed the performance of our estimator in the case where causal effect sizes are identical between populations. We find that even when causal effect sizes are simulated to be identical between populations, the marginal effect sizes at trait-associated SNPs differ enough that our model remains identifiable and we estimate values of *δ* close to 0 (Figure S6). We additionally investigate this in empirical data by estimating *δ* from two subsets of European Americans between which we expect causal effect sizes to be identical. For both gene expression and LDL-C, we estimate values of *δ* close to 0, demonstrating that our estimator 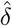 has the desired behavior when causal effect sizes are identical between populations (Figure S7).

Finally, we used our model to test whether causal variants have the same effect size in African Americans and European Americans. For gene expression, 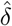 is significantly different from zero, with a maximum likelihood estimate (MLE) of 0.47 and a 95% CI of (0.39, 0.53) (Figure 3B). For LDL-C, we estimate a similar MLE of 0.46 with a 95% CI of (−0.06, 0.87) (Figure 3C). Moreover, we find that the term containing *δ* contributes modestly to phenotypic variance: 0.01% for gene expression, 0.01% LDL-C. Thus, our results indicate that SNPs in regions of European ancestry in African Americans and European Americans have different marginal effect sizes, suggesting that causal variant effect sizes differ between populations because they are modified by the genome or environment, providing evidence for gene-by-gene or gene-by-environment interactions.

**Figure 3:**
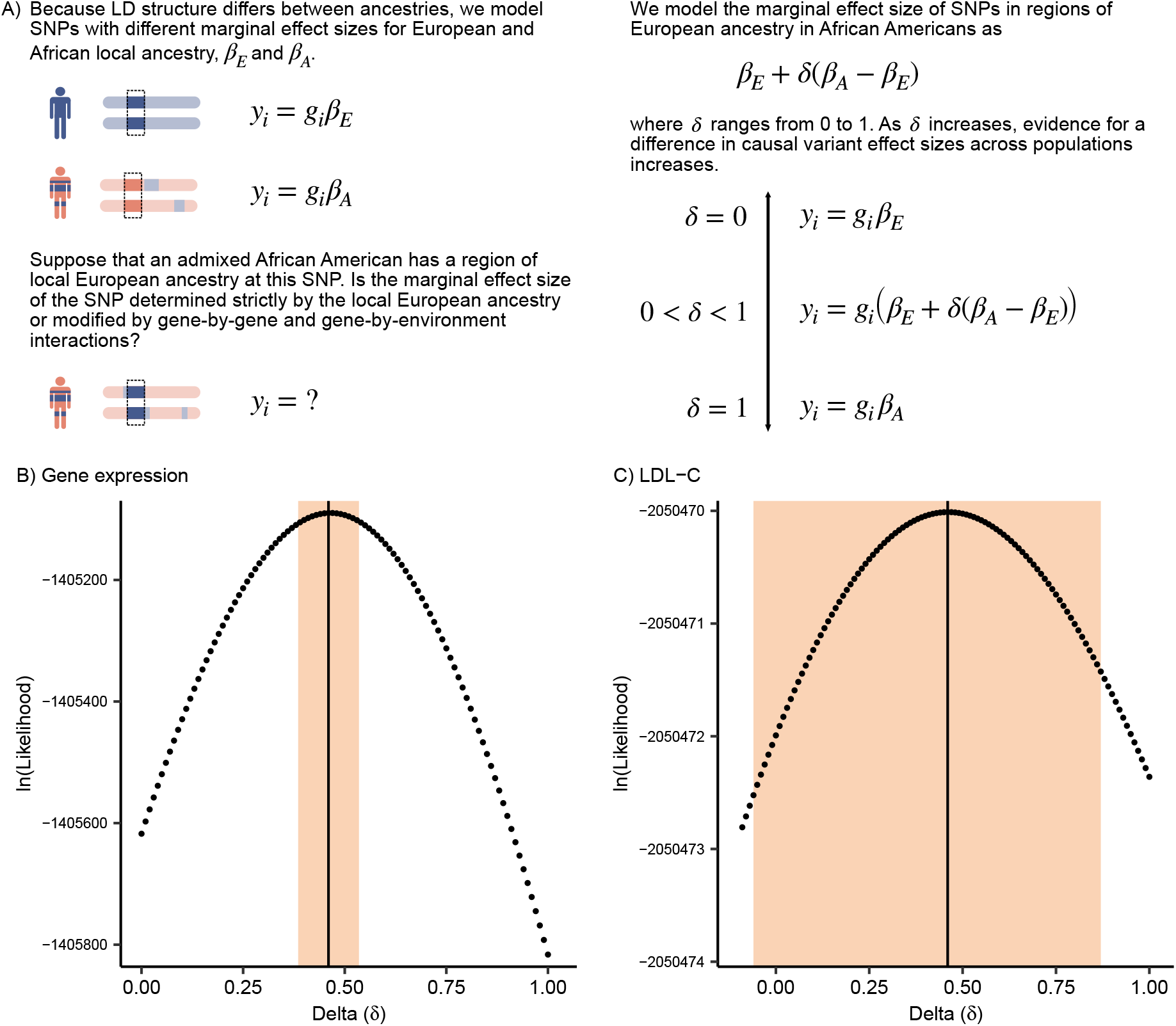
Testing for genetic interactions. **A)** We looked for evidence of genetic interactions by testing for differences in causal variant effect sizes between African Americans and European Americans. The parameter *δ* measures the extent to which the marginal effect sizes of SNPs in regions of European ancestry in African Americans differ from those in European Americans. **B, C)** Likelihood surface for *δ*. Maximum likelihood estimates and 95% bootstrap CI are 0.47 (0.39, 0.53) for gene expression and 0.46 (−0.06, 0.87) for LDL-C. We denote the MLE and 95% bootstrap CI with the vertical line and shaded region, respectively.

## Discussion

We developed a model in which we introduce the parameter *δ* to test for the existence of genetic interactions. Specifically, we leveraged regions of European ancestry shared between African Americans and European Americans to compare marginal effect sizes of trait-associated SNPs in a manner unbiased by LD structure. We applied our model to two traits, gene expression in MESA and LDL-C in MVP. For gene expression, we observe that 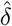 is significantly different from zero, implying that causal variant effect sizes differ between African Americans and European Americans. For LDL-C, we obtain a MLE for *δ* that is similar to that from gene expression but not significantly different from zero. These observed differences in causal variant effect sizes between populations must be due to unmodeled gene-by-gene or gene-by-environment interactions. Our observation that causal variant effect sizes differ between populations is also relevant to previous work on quantifying cross-population genetic correlations^13,14^. There is no straightforward analytical relationship between our parameter *δ* and genetic correlation, but our results are intuitively consistent with a cross-population genetic correlation less than one.

Though we observe that causal variant effect sizes significantly differ between populations, we also find that the inclusion of the *δ* term in the model does not substantially increase the amount of phenotypic variance explained. This apparent discrepancy can be resolved by noting that we evaluate model performance on the full dataset of African Americans and European Americans, but the *δ* term will only improve the modeling of effect sizes in regions of European ancestry in African Americans, which only represents about 10% of the full dataset.

Our results have implications for modeling complex trait phenotypes with polygenic scores (PGS). We find that trait-associated SNPs ascertained in Europeans have attenuated effect sizes in African Americans, which is consistent with European-ascertained SNPs tagging causal variants poorly in African ancestry. Thus, our findings corroborate earlier work demonstrating that differences in LD structure contribute to poor PGS portability, reiterating that a PGS will perform best when constructed from a population with similar LD structure^12,37–40^. Moreover, our findings imply the existence of genetic interactions, which challenges the assumption of additivity made by the statistical genetic models underpinning PGS. This suggests that genetic interactions could contribute to poor PGS portability, though it remains unclear to what extent they may do so.

Future directions include applying our model to additional traits. The larger confidence interval we observe for LDL-C is likely due to differences in statistical power between the two traits. Though we used significantly associated SNPs for both traits, many fewer SNPs were used in LDL-C analyses (122 SNPs) than in gene expression analyses (4,236 SNPs). Moreover, trait-associated SNPs were ascertained within the same dataset (MESA) for gene expression but were ascertained from an external dataset (UK Biobank) for LDL-C. This should not bias the estimation of *δ* but may mean that trait-associated SNPs capture a larger proportion of phenotypic variance for gene expression relative to LDL-C. Thus, by applying our model to additional traits, such as those with thousands of associated SNPs, we could gain further insights into the role of genetic interactions in complex traits. Another area of investigation includes adapting our model to understand how the magnitude of genetic interactions varies across SNPs or individuals. We only estimate one parameter *δ* from all trait-associated SNPs in order to maximize power, but by understanding how *δ* varies with certain functional genomic properties of SNPs or with individuals’ ancestry, we could begin to untangle the contributions of gene-by-gene versus gene-by-environment interactions.

In summary, we find evidence for genetic interactions by testing for differences in causal variant effect sizes between populations. This analysis is motivated by the assumption that the African American and European American individuals in our datasets have sufficiently different genetic and environmental backgrounds such that the existence of gene-by-gene or gene-by-environment interactions will produce modest differences in causal variant effect sizes. However, we reiterate others’ findings that there is a great deal of genetic and environmental heterogeneity within human populations^41,42,40^. Thus, it is worth noting that if causal variant effect sizes can be modified by gene-by-gene or gene-by-environment interactions, it follows that causal variant effect sizes will differ not only between populations, but also between individuals within a population. Ultimately, our results give insight into the importance of genetic interactions in human complex traits.

## Supporting information

Supplemental Information - MVP authors

## Ethics

Human subjects: This research has been conducted using the Multi-Ethnic Study of Atherosclerosis (MESA) dataset, the Million Veteran Program (MVP) dataset, and the UK Biobank dataset. The MESA dataset was obtained under TOPMed application number 10194, ”Investigating cross-population portability of variant effect sizes”. All MESA participants provided written informed consent. The MVP dataset was obtained under MVP application number 200229, ”Genetics of Cardiometabolic Diseases in the VA population”. All MVP participants provided written informed consent, and the study protocol was approved by the Veterans Affairs Central Institutional Review Board. The UK Biobank dataset was obtained under application number 24983, ”Generating effective therapeutic hypotheses from genomic and hospital linkage data”. All participants of UK Biobank provided written informed consent.

## Data and Code Availability

The code generated during this study is available on GitHub (https://github.com/roshnipatel/LocalAncestry; https://github.com/roshnipatel/eQTLs). The Multi-Ethnic Study of Atherosclerosis (MESA) and the Million Veteran Program (MVP) datasets are available via dbGAP with study accessions phs000209.v13.p3 and phs001672.v6.p1, respectively.

## Declaration of Interests

The authors declare no competing interests.

## Acknowledgments

We thank Matthew Aguirre, Margaret Antonio, Nicole Ersaro, Nicole Gay, Arbel Harpak, Kangcheng Hou, Stephen Montgomery, Bogdan Pasaniuc, Molly Przeworski, and additional members of the Pritchard lab for helpful comments and/or technical advice on this project. Research reported in this publication was supported by the National Center For Advancing Translational Sciences of the National Institutes of Health under Award Number UL1TR003142. The content is solely the responsibility of the authors and does not necessarily represent the official views of the National Institutes of Health. This work was supported by NIH grants R01HG008140, U01HG009431, R01HL142015, and R01HG011432. Molecular data for the Trans-Omics in Precision Medicine (TOPMed) program was supported by the National Heart, Lung and Blood Institute (NHLBI). Multi-Ethnic Study of Atherosclerosis (MESA)” (phs001416.v1.p1) was performed at the Broad Institute of MIT and Harvard (3U54HG003067-13S1). Centralized read mapping and genotype calling, along with variant quality metrics and filtering were provided by the TOPMed Informatics Research Center (3R01HL-117626-02S1, contract HHSN268201800002I) (Broad RNA Seq, Proteomics HHSN268201600034I, UW RNA Seq HHSN268201600032I, USC DNA Methylation HHSN268201600034I, Broad Metabolomics HHSN268201600038I). Phenotype harmonization, data management, sample-identity QC, and general study coordination, were provided by the TOPMed Data Coordinating Center (3R01HL-120393; U01HL-120393; contract HHSN268180001I). The MESA project is conducted and supported by the National Heart, Lung, and Blood Institute (NHLBI) in collaboration with MESA investigators. Support for MESA is provided by contracts 75N92020D00001, HHSN268201500003I, N01-HC-95159, 75N92020D00005, N01-HC-95160, 75N92020D00002, N01-HC-95161, 75N92020D00003, N01-HC-95162, 75N92020D00006, N01-HC-95163, 75N92020D00004, N01-HC-95164, 75N92020D00007, N01-HC-95165, N01-HC-95166, N01-HC-95167, N01-HC-95168, N01-HC-95169, UL1-TR-000040, UL1-TR-001079, UL1-TR-001420. Also supported in part by the National Center for Advancing Translational Sciences, CTSI grant UL1TR001881, and the National Institute of Diabetes and Digestive and Kidney Disease Diabetes Research Center (DRC) grant DK063491 to the Southern California Diabetes Endocrinology Research Center. Infrastructure for the CHARGE Consortium is supported in part by the National Heart, Lung, and Blood Institute (NHLBI) grant R01HL105756. This research is based on data from the Million Veteran Program, Office of Research and Development, Veterans Health Administration, and was supported by award I01BX003362. This publication does not represent the views of the Department of Veteran Affairs or the United States Government.

## Supplementary Items

**Figure S1:**
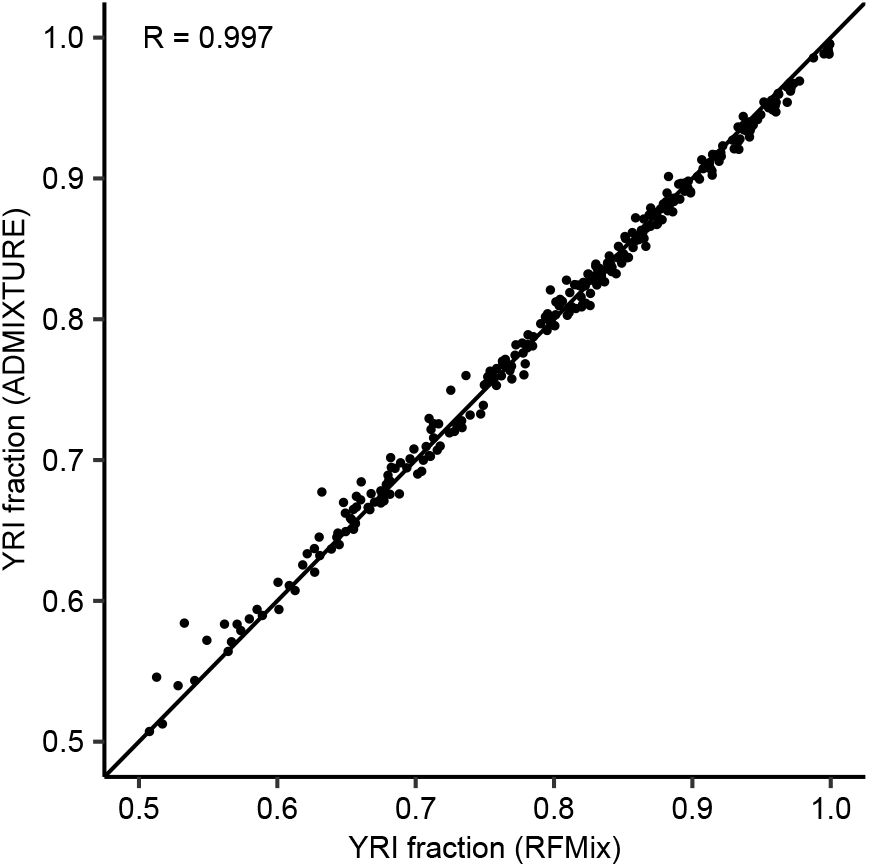
Comparison of ancestry inference methods. We observe a strong correlation between RFMix and ADMIXTURE estimates of global African ancestry fraction for African American individuals in MESA.

**Figure S2:**
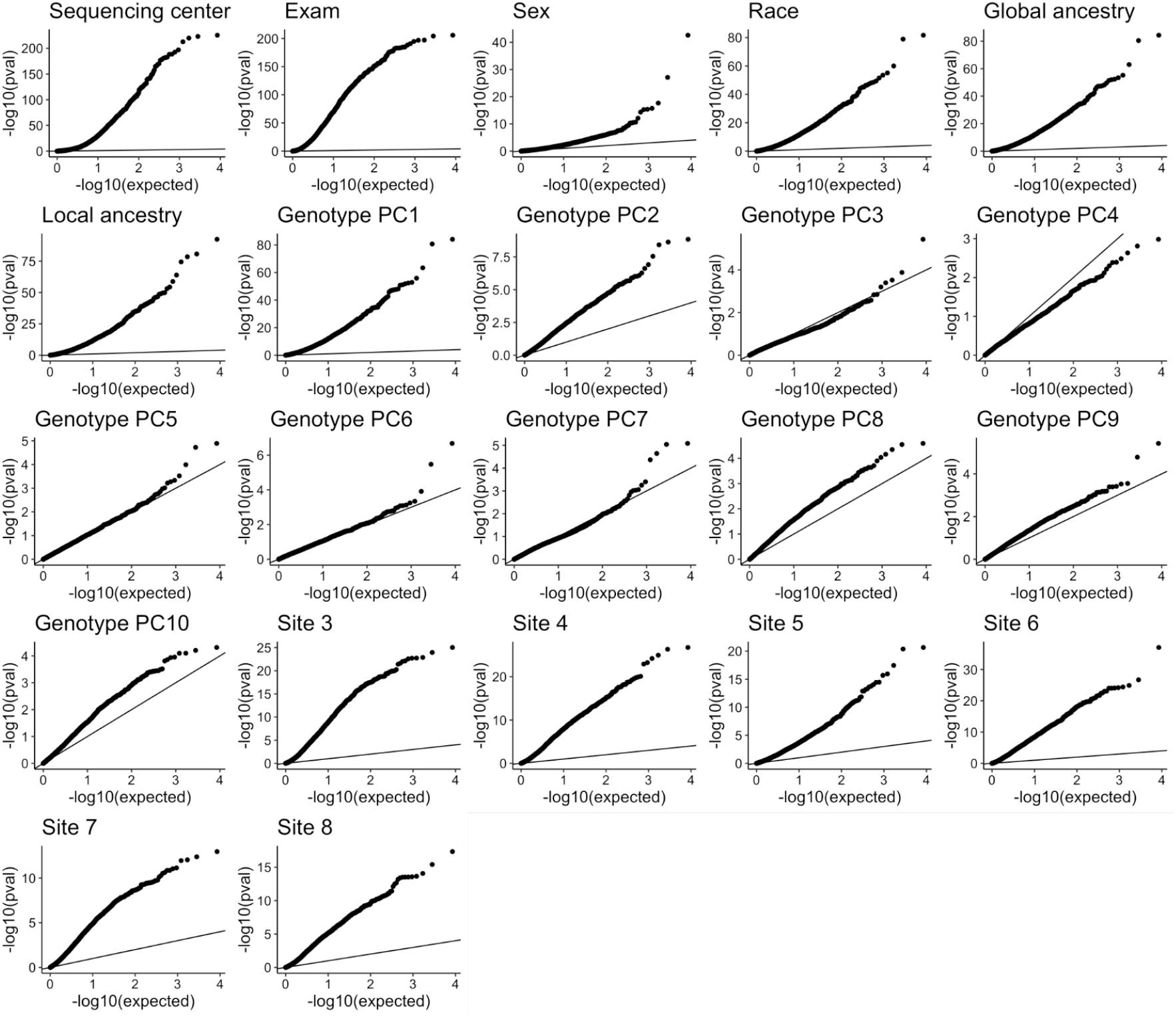
Association between gene expression phenotypes and covariates. We tested for statistical association between phenotypes of all 4,236 significant genes and 22 covariates, including 2 batch covariates (sequencing center and time of exam), sex, race, global and local ancestry, 10 genotype principal components (PCs), and 6 covariates corresponding to a one-hot encoding of recruitment site. We show the resulting QQ plots of association p-values, demonstrating that expression phenotypes are significantly associated with sequencing center, time of exam, sex, race, global and local ancestry, recruitment site, and the first two PCs.

**Figure S3:**
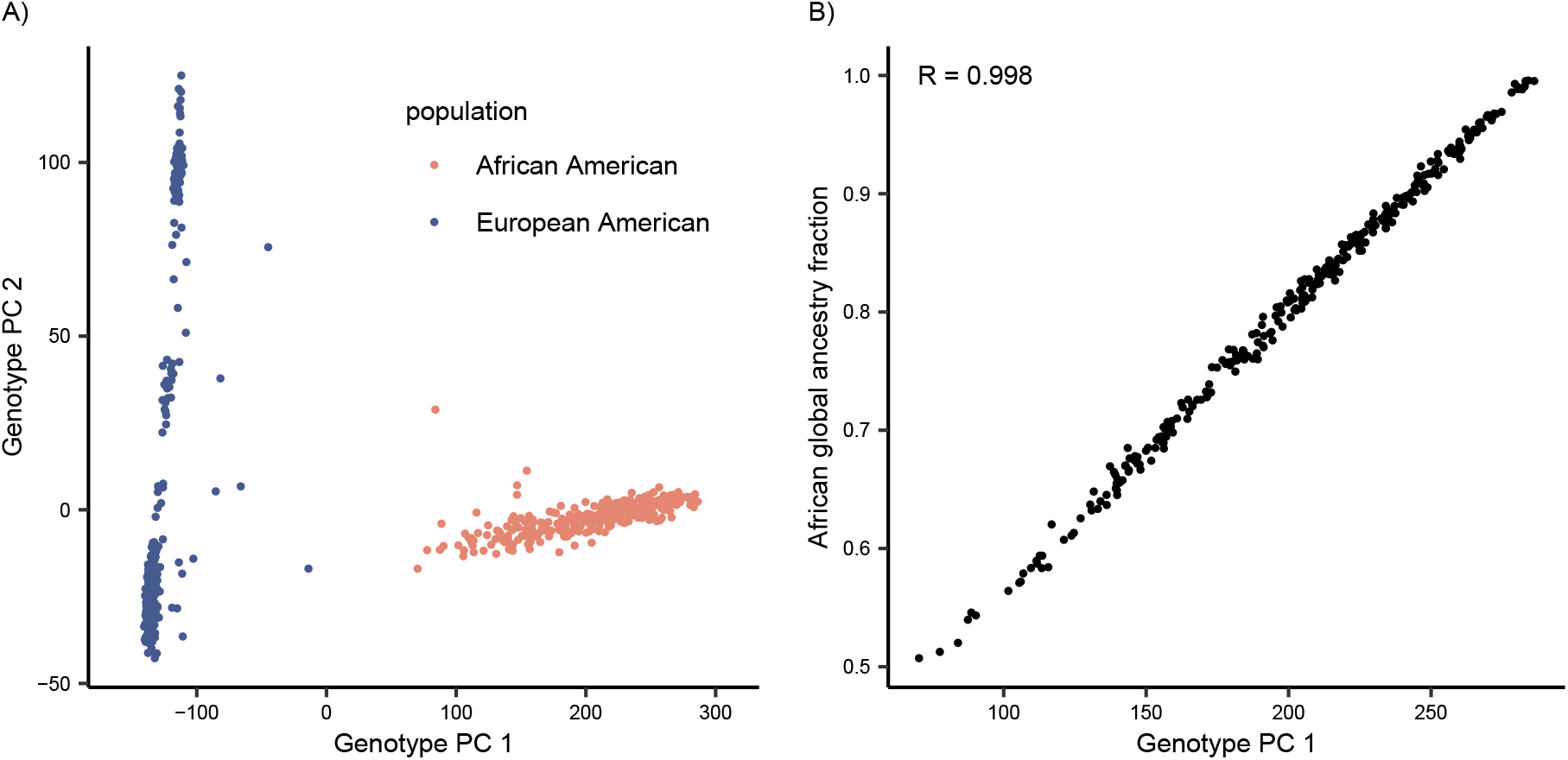
Principal components analysis of MESA genotypes. **A)** We computed principal components from the genotypes of 296 African Americans and 482 European Americans in MESA. The first genotype PC stratifies the African Americans and the second genotype PC stratifies the European Americans. **B)** Within African Americans, the first genotype PC is highly correlated with African global ancestry fraction.

**Figure S4:**
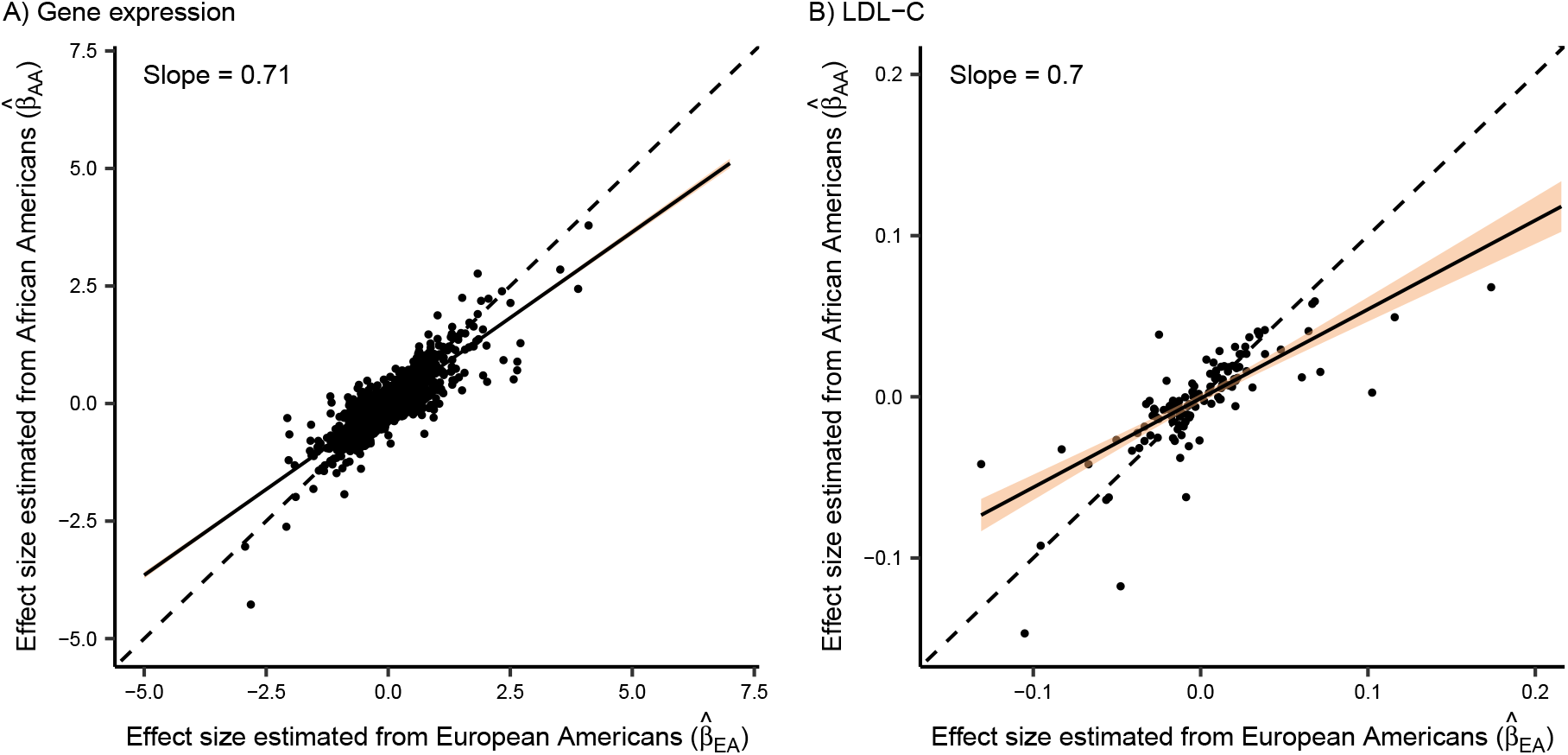
Ordinary least squares regression of *β*_*AA*_ on *β*_*EA*_ for **A)** gene expression and **B)** LDL-C. We represent the 95% CI with the shaded region.

**Figure S5:**
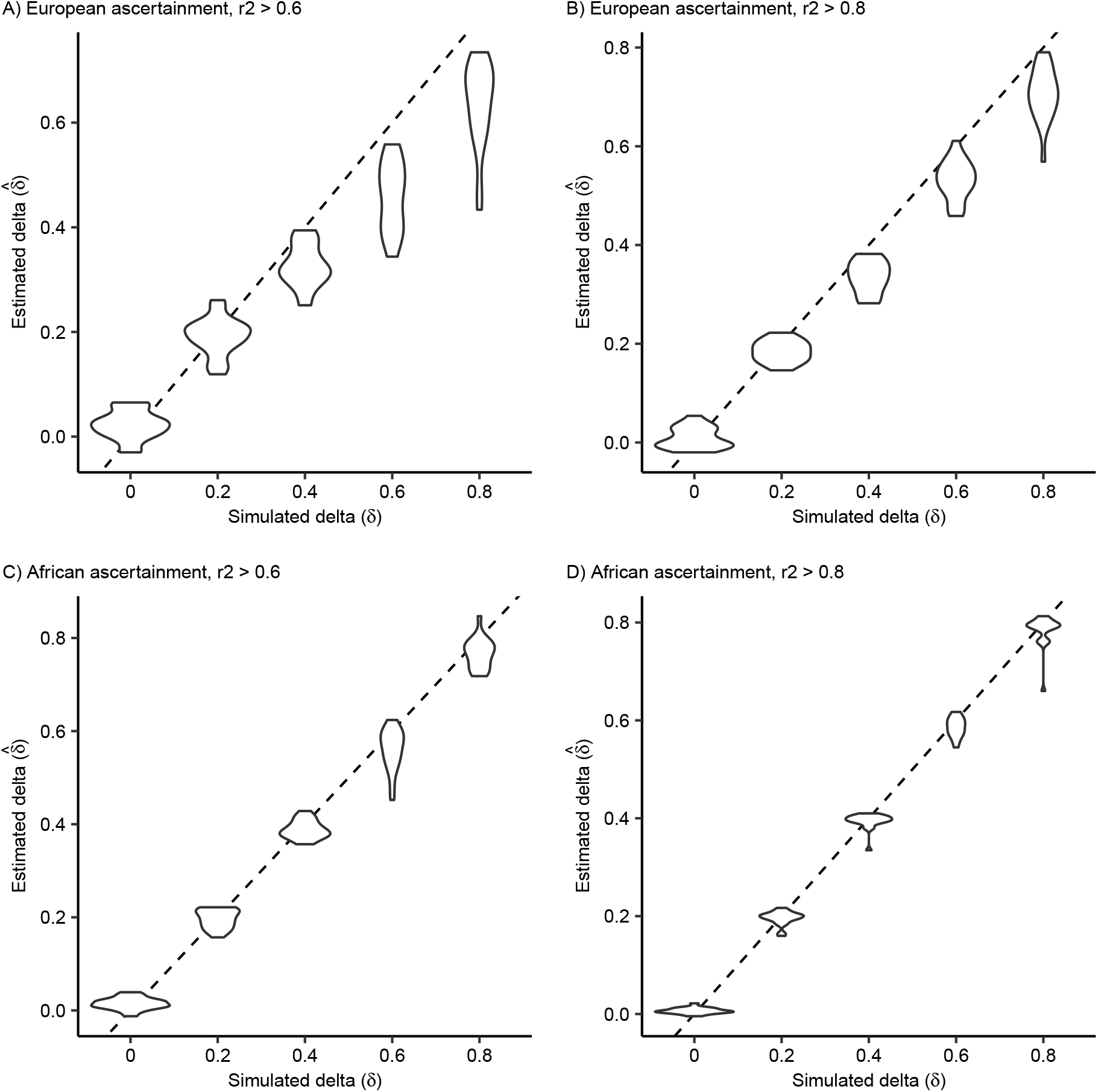
Estimates of *δ* from simulations where causal variant effect sizes are allowed to differ between populations. We simulated ascertainment in both European and African ancestries and required that the squared correlation between the causal SNP and the tag SNP was either greater than 0.6 or 0.8.

**Figure S6:**
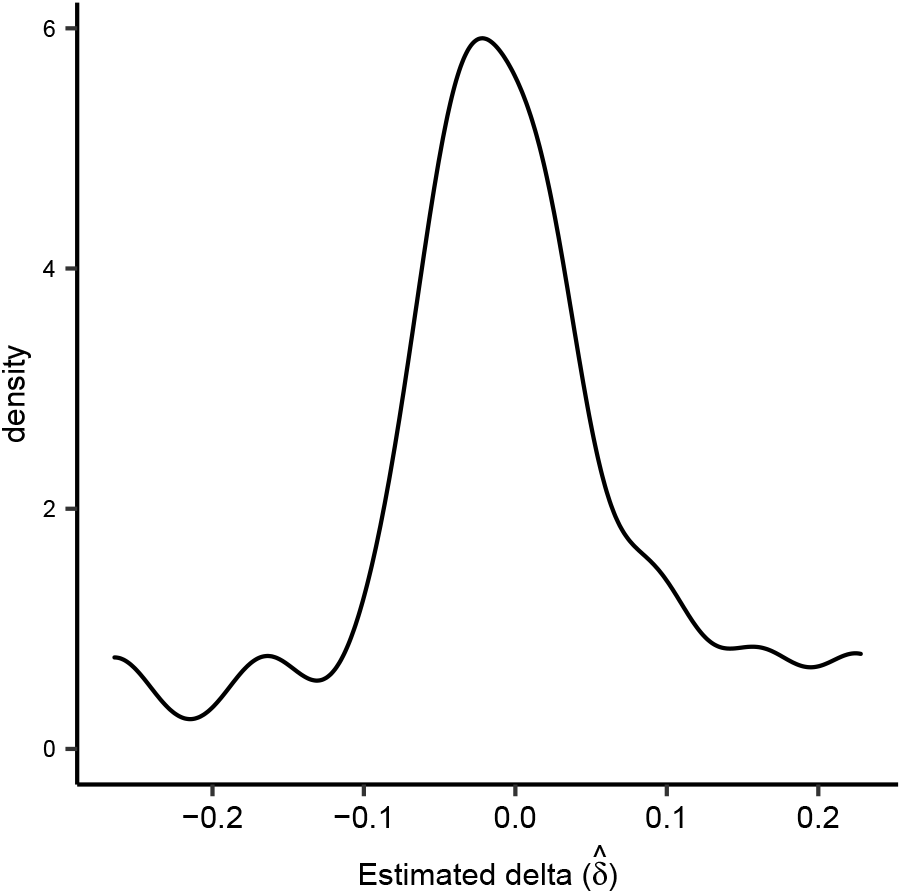
Estimates of *δ* from ten simulations where causal variant effect sizes are identical between populations.

**Figure S7:**
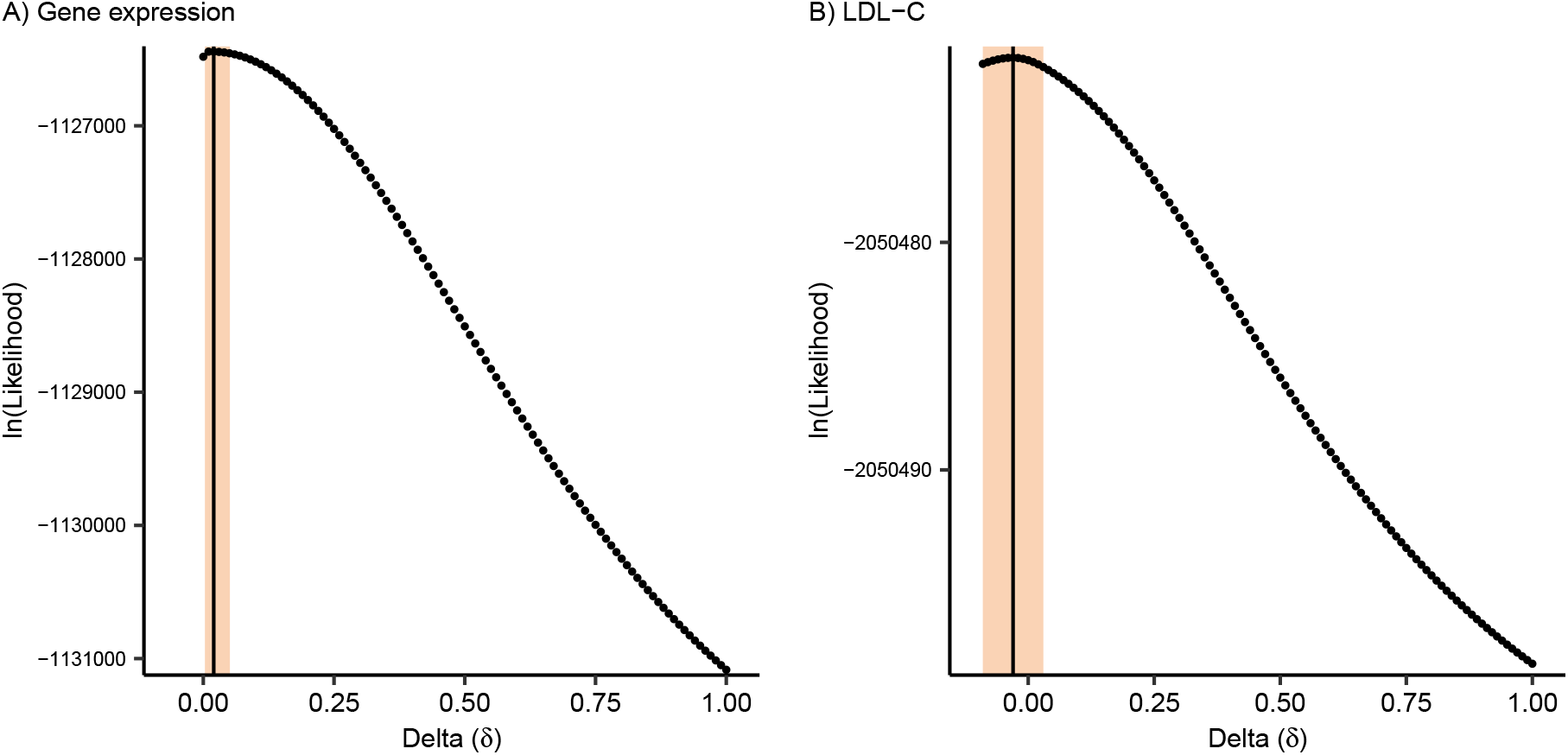
Likelihood surface for *δ* when comparing causal variant effect sizes between two subsets of European Americans. Maximum likelihood estimates and 95% bootstrap CI are **A)** 0.008 (0.003, 0.05) for gene expression and **B)** -0.03 (−0.09, 0.03) for LDL-C. We denote the MLE and 95% bootstrap CI with the vertical line and shaded region, respectively.

## Notes

### Competing Interest Statement

The authors have declared no competing interest.

### Summary of Updates

The title and Introduction have been revised to better capture the main contributions of our paper; the Results have been revised to better clarify the assumptions underlying our statistical model.

