## Supplemental Information - MVP authors for "Genetic interactions drive heterogeneity in causal variant effect sizes for gene expression and complex traits"

**Consortium Authors: VA Million Veteran Program**
**MVP Executive Committee**

- Co-Chair: J. Michael Gaziano, M.D., M.P.H.

VA Boston Healthcare System, 150 S. Huntington Avenue, Boston, MA 02130

- Co-Chair: Sumitra Muralidhar, Ph.D.

US Department of Veterans Affairs, 810 Vermont Avenue NW, Washington, DC 20420

- Rachel Ramoni, D.M.D., Sc.D., Chief VA Research and Development Officer

US Department of Veterans Affairs, 810 Vermont Avenue NW, Washington, DC 20420

- Jean Beckham, Ph.D.

Durham VA Medical Center, 508 Fulton Street, Durham, NC 27705

- Kyong-Mi Chang, M.D.

Philadelphia VA Medical Center, 3900 Woodland Avenue, Philadelphia, PA 19104

- Christopher J. O'Donnell, M.D., M.P.H.

VA Boston Healthcare System, 150 S. Huntington Avenue, Boston, MA 02130

- Philip S. Tsao, Ph.D.

VA Palo Alto Health Care System, 3801 Miranda Avenue, Palo Alto, CA 94304

- James Breeling, M.D., Ex-Officio

US Department of Veterans Affairs, 810 Vermont Avenue NW, Washington, DC 20420

- Grant Huang, Ph.D., Ex-Officio

US Department of Veterans Affairs, 810 Vermont Avenue NW, Washington, DC 20420

- JP Casas Romero, M.D., Ph.D., Ex-Officio

VA Boston Healthcare System, 150 S. Huntington Avenue, Boston, MA 02130

**MVP Program Office**

- Sumitra Muralidhar, Ph.D.

US Department of Veterans Affairs, 810 Vermont Avenue NW, Washington, DC 20420

- Jennifer Moser, Ph.D.

US Department of Veterans Affairs, 810 Vermont Avenue NW, Washington, DC 20420

**MVP Recruitment/Enrollment**

- Recruitment/Enrollment Director/Deputy Director, Boston – Stacey B. Whitbourne, Ph.D.; Jessica V. Brewer, M.P.H.

VA Boston Healthcare System, 150 S. Huntington Avenue, Boston, MA 02130

- MVP Coordinating Centers
    - Clinical Epidemiology Research Center (CERC), West Haven – Mihaela Aslan, Ph.D.

      West Haven VA Medical Center, 950 Campbell Avenue, West Haven, CT 06516
    - Cooperative Studies Program Clinical Research Pharmacy Coordinating Center, Albuquerque – Todd Connor, Pharm.D.; Dean P. Argyres, B.S., M.S.

      New Mexico VA Health Care System, 1501 San Pedro Drive SE, Albuquerque, NM 87108
    - Genomics Coordinating Center, Palo Alto – Philip S. Tsao, Ph.D.

      VA Palo Alto Health Care System, 3801 Miranda Avenue, Palo Alto, CA 94304
    - MVP Boston Coordinating Center, Boston - J. Michael Gaziano, M.D., M.P.H.

      VA Boston Healthcare System, 150 S. Huntington Avenue, Boston, MA 02130
    - MVP Information Center, Canandaigua – Brady Stephens, M.S.

      Canandaigua VA Medical Center, 400 Fort Hill Avenue, Canandaigua, NY 14424
- VA Central Biorepository, Boston – Mary T. Brophy M.D., M.P.H.; Donald E. Humphries, Ph.D.; Luis E. Selva, Ph.D.

VA Boston Healthcare System, 150 S. Huntington Avenue, Boston, MA 02130

- MVP Informatics, Boston – Nhan Do, M.D.; Shahpoor Shayan

VA Boston Healthcare System, 150 S. Huntington Avenue, Boston, MA 02130

- MVP Data Operations/Analytics, Boston – Kelly Cho, Ph.D.

VA Boston Healthcare System, 150 S. Huntington Avenue, Boston, MA 02130

**MVP Science**

- Science Operations – Christopher J. O'Donnell, M.D., M.P.H.

VA Boston Healthcare System, 150 S. Huntington Avenue, Boston, MA 02130

- Genomics Core - Christopher J. O'Donnell, M.D., M.P.H.; Saiju Pyarajan Ph.D.

VA Boston Healthcare System, 150 S. Huntington Avenue, Boston, MA 02130

Philip S. Tsao, Ph.D.

VA Palo Alto Health Care System, 3801 Miranda Avenue, Palo Alto, CA 94304

- Phenomics Core- Kelly Cho, M.P.H, Ph.D.

VA Boston Healthcare System, 150 S. Huntington Avenue, Boston, MA 02130

- Data and Computational Sciences – Saiju Pyarajan, Ph.D.

VA Boston Healthcare System, 150 S. Huntington Avenue, Boston, MA 02130

- Statistical Genetics – Elizabeth Hauser, Ph.D.

Durham VA Medical Center, 508 Fulton Street, Durham, NC 27705

Yan Sun, Ph.D.

Atlanta VA Medical Center, 1670 Clairmont Road, Decatur, GA 30033

Hongyu Zhao, Ph.D.

West Haven VA Medical Center, 950 Campbell Avenue, West Haven, CT 06516

**Current MVP Local Site Investigators**

- Atlanta VA Medical Center (Peter Wilson, M.D.)

1670 Clairmont Road, Decatur, GA 30033

- Bay Pines VA Healthcare System (Rachel McArdle, Ph.D.)

10,000 Bay Pines Blvd Bay Pines, FL 33744

- Birmingham VA Medical Center (Louis Dellitalia, M.D.)

700 S. 19th Street, Birmingham AL 35233

- Central Western Massachusetts Healthcare System (Kristin Mattocks, Ph.D., M.P.H.)

421 North Main Street, Leeds, MA 01053

- Cincinnati VA Medical Center (John Harley, M.D., Ph.D.)

3200 Vine Street, Cincinnati, OH 45220

- Clement J. Zablocki VA Medical Center (Jeffrey Whittle, M.D., M.P.H.)

5000 West National Avenue, Milwaukee, WI 53295

- VA Northeast Ohio Healthcare System (Frank Jacono, M.D.)

10701 East Boulevard, Cleveland, OH 44106

- Durham VA Medical Center (Jean Beckham, Ph.D.)

508 Fulton Street, Durham, NC 27705

- Edith Nourse Rogers Memorial Veterans Hospital (John Wells., Ph.D.)

200 Springs Road, Bedford, MA 01730

- Edward Hines, Jr. VA Medical Center (Salvador Gutierrez, M.D.)

5000 South 5th Avenue, Hines, IL 60141

- Veterans Health Care System of the Ozarks (Gretchen Gibson, D.D.S., M.P.H.)

1100 North College Avenue, Fayetteville, AR 72703

- Fargo VA Health Care System (Kimberly Hammer, Ph.D.)

2101 N. Elm, Fargo, ND 58102

- VA Health Care Upstate New York (Laurence Kaminsky, Ph.D.)

113 Holland Avenue, Albany, NY 12208

- New Mexico VA Health Care System (Gerardo Villareal, M.D.)

1501 San Pedro Drive, S.E. Albuquerque, NM 87108

- VA Boston Healthcare System (Scott Kinlay, M.B.B.S., Ph.D.)

150 S. Huntington Avenue, Boston, MA 02130

- VA Western New York Healthcare System (Junzhe Xu, M.D.)

3495 Bailey Avenue, Buffalo, NY 14215-1199

- Ralph H. Johnson VA Medical Center (Mark Hamner, M.D.)

109 Bee Street, Mental Health Research, Charleston, SC 29401

- Columbia VA Health Care System (Roy Mathew, M.D.)

6439 Garners Ferry Road, Columbia, SC 29209

- VA North Texas Health Care System (Sujata Bhushan, M.D.)

4500 S. Lancaster Road, Dallas, TX 75216

- Hampton VA Medical Center (Pran Iruvanti, D.O., Ph.D.)

100 Emancipation Drive, Hampton, VA 23667

- Richmond VA Medical Center (Michael Godschalk, M.D.)

1201 Broad Rock Blvd., Richmond, VA 23249

- Iowa City VA Health Care System (Zuhair Ballas, M.D.)

601 Highway 6 West, Iowa City, IA 52246-2208

- Eastern Oklahoma VA Health Care System (Douglas Ivins, M.D.)

1011 Honor Heights Drive, Muskogee, OK 74401

- James A. Haley Veterans' Hospital (Stephen Mastorides, M.D.)

13000 Bruce B. Downs Blvd, Tampa, FL 33612

- James H. Quillen VA Medical Center (Jonathan Moorman, M.D., Ph.D.)

Corner of Lamont & Veterans Way, Mountain Home, TN 37684

- John D. Dingell VA Medical Center (Saib Gappy, M.D.)

4646 John R Street, Detroit, MI 48201

- Louisville VA Medical Center (Jon Klein, M.D., Ph.D.)

800 Zorn Avenue, Louisville, KY 40206

- Manchester VA Medical Center (Nora Ratcliffe, M.D.)

718 Smyth Road, Manchester, NH 03104

- Miami VA Health Care System (Hermes Florez, M.D., Ph.D.)

1201 NW 16th Street, 11 GRC, Miami FL 33125

- Michael E. DeBakey VA Medical Center (Olaoluwa Okusaga, M.D.)

2002 Holcombe Blvd, Houston, TX 77030

- Minneapolis VA Health Care System (Maureen Murdoch, M.D., M.P.H.)

One Veterans Drive, Minneapolis, MN 55417

- N. FL/S. GA Veterans Health System (Peruvemba Sriram, M.D.)

1601 SW Archer Road, Gainesville, FL 32608

- Northport VA Medical Center (Shing Shing Yeh, Ph.D., M.D.)

79 Middleville Road, Northport, NY 11768

- Overton Brooks VA Medical Center (Neeraj Tandon, M.D.)

510 East Stoner Ave, Shreveport, LA 71101

- Philadelphia VA Medical Center (Darshana Jhala, M.D.)

3900 Woodland Avenue, Philadelphia, PA 19104

- Phoenix VA Health Care System (Samuel Aguayo, M.D.)

650 E. Indian School Road, Phoenix, AZ 85012

- Portland VA Medical Center (David Cohen, M.D.)

3710 SW U.S. Veterans Hospital Road, Portland, OR 97239

- Providence VA Medical Center (Satish Sharma, M.D.)

830 Chalkstone Avenue, Providence, RI 02908

- Richard Roudebush VA Medical Center (Suthat Liangpunsakul, M.D., M.P.H.)

1481 West 10th Street, Indianapolis, IN 46202

- Salem VA Medical Center (Kris Ann Oursler, M.D.)

1970 Roanoke Blvd, Salem, VA 24153

- San Francisco VA Health Care System (Mary Whooley, M.D.)

4150 Clement Street, San Francisco, CA 94121

- South Texas Veterans Health Care System (Sunil Ahuja, M.D.)

7400 Merton Minter Boulevard, San Antonio, TX 78229

- Southeast Louisiana Veterans Health Care System (Joseph Constans, Ph.D.)

2400 Canal Street, New Orleans, LA 70119

- Southern Arizona VA Health Care System (Paul Meyer, M.D., Ph.D.)

3601 S 6th Avenue, Tucson, AZ 85723

- Sioux Falls VA Health Care System (Jennifer Greco, M.D.)

2501 W 22nd Street, Sioux Falls, SD 57105

- St. Louis VA Health Care System (Michael Rauchman, M.D.)

915 North Grand Blvd, St. Louis, MO 63106

- Syracuse VA Medical Center (Richard Servatius, Ph.D.)

800 Irving Avenue, Syracuse, NY 13210

- VA Eastern Kansas Health Care System (Melinda Gaddy, Ph.D.)

4101 S 4th Street Trafficway, Leavenworth, KS 66048

- VA Greater Los Angeles Health Care System (Agnes Wallbom, M.D., M.S.)

11301 Wilshire Blvd, Los Angeles, CA 90073

- VA Long Beach Healthcare System (Timothy Morgan, M.D.)

5901 East 7th Street Long Beach, CA 90822

- VA Maine Healthcare System (Todd Stapley, D.O.)

1 VA Center, Augusta, ME 04330

- VA New York Harbor Healthcare System (Scott Sherman, M.D., M.P.H.)

423 East 23rd Street, New York, NY 10010

- VA Pacific Islands Health Care System (George Ross, M.D.)

459 Patterson Rd, Honolulu, HI 96819

- VA Palo Alto Health Care System (Philip Tsao, Ph.D.)

3801 Miranda Avenue, Palo Alto, CA 94304-1290

- VA Pittsburgh Health Care System (Patrick Strollo, Jr., M.D.)

University Drive, Pittsburgh, PA 15240

- VA Puget Sound Health Care System (Edward Boyko, M.D.)

1660 S. Columbian Way, Seattle, WA 98108-1597

- VA Salt Lake City Health Care System (Laurence Meyer, M.D., Ph.D.)

500 Foothill Drive, Salt Lake City, UT 84148

- VA San Diego Healthcare System (Samir Gupta, M.D., M.S.C.S.)

3350 La Jolla Village Drive, San Diego, CA 92161

- VA Sierra Nevada Health Care System (Mostaqul Huq, Pharm.D., Ph.D.)

975 Kirman Avenue, Reno, NV 89502

- VA Southern Nevada Healthcare System (Joseph Fayad, M.D.)

6900 North Pecos Road, North Las Vegas, NV 89086

- VA Tennessee Valley Healthcare System (Adriana Hung, M.D., M.P.H.)

1310 24th Avenue, South Nashville, TN 37212

- Washington DC VA Medical Center (Jack Lichy, M.D., Ph.D.)

50 Irving St, Washington, D. C. 20422

- W.G. (Bill) Hefner VA Medical Center (Robin Hurley, M.D.)

1601 Brenner Ave, Salisbury, NC 28144

- White River Junction VA Medical Center (Brooks Robey, M.D.)

163 Veterans Drive, White River Junction, VT 05009

- William S. Middleton Memorial Veterans Hospital (Robert Striker, M.D., Ph.D.)

2500 Overlook Terrace, Madison, WI 53705
